# Direct cochlear recordings in humans show a theta rhythmic modulation of the auditory nerve by selective attention

**DOI:** 10.1101/2021.03.01.433316

**Authors:** Quirin Gehmacher, Patrick Reisinger, Thomas Hartmann, Thomas Keintzel, Sebastian Rösch, Konrad Schwarz, Nathan Weisz

## Abstract

The architecture of the efferent auditory system enables prioritization of strongly overlapping spatiotemporal cochlear activation patterns elicited by relevant and irrelevant inputs. So far, attempts at finding such attentional modulations of cochlear activity delivered indirect insights in humans or required direct recordings in animals. The extent to which spiral ganglion cells forming the human auditory nerve are sensitive to selective attention remains largely unknown. We investigated this question by testing the effects of attending to either the auditory or visual modality in human cochlear implant (CI) users (3 female, 13 male). Auditory nerve activity was directly recorded with standard CIs during a silent (anticipatory) cue-target interval. When attending the upcoming auditory input, ongoing auditory nerve activity within the theta range (5-8 Hz) was enhanced. Crucially, using the broadband signal (4-25 Hz), a classifier was even able to decode the attended modality from single-trial data. Follow-up analysis showed that the effect was not driven by a narrow frequency in particular. Using direct cochlear recordings from deaf individuals, our findings suggest that cochlear spiral ganglion cells are sensitive to top-down attentional modulations. Given the putatively broad hair-cell degeneration of these individuals, the effects are likely mediated by alternative efferent pathways as compared to previous studies using otoacoustic emissions. Successful classification of single-trial data could additionally have a significant impact on future closed-loop CI developments that incorporate real-time optimization of CI parameters based on the current mental state of the user.

**Significance Statement:** The efferent auditory system in principle allows top-down modulation of auditory nerve activity, however evidence for this is lacking in humans. Using cochlear recordings in participants performing an audiovisual attention task, we show that ongoing auditory nerve activity in the silent cue-target period is directly modulated by selective attention. Specifically, ongoing auditory nerve activity is enhanced within the theta range when attending upcoming auditory input. Furthermore, over a broader frequency range, the attended modality can be decoded from single-trial data. Demonstrating this direct top-down influence on auditory nerve activity substantially extends previous works that focus on outer hair cell activity. Generally, our work could promote the use of standard cochlear implant electrodes to study cognitive neuroscientific questions.

## Introduction

Attention describes a process by which sensory information can be prioritized. For all sensory modalities common spatiotemporal cortical activity patterns have been reported, suggesting modality independent mechanisms to select or ignore features by alterations of oscillatory activity in the alpha (Frey et al., 2014; Mazaheri et al., 2014; Weise et al., 2016) and beta band (Buschman and Miller, 2007; Iversen et al., 2009; Lee et al., 2013). As the auditory system comprises a unique complex subcortical network (Terreros and Delano, 2015; Elgueda and Delano, 2020), cochlear activity can in principle be altered by top-down signals from the auditory cortex via only one extra relay through the superior olivary complex (SOC). However, studying peripheral attentional mechanisms requires special recording and analysis techniques (Elgueda and Delano, 2020) and has therefore been rarely investigated.

Noninvasively, evidence in humans comes from studies on otoacoustic emissions (OAEs), sounds that are generated by outer hair cell (OHC) activity in the cochlea. OHCs are modulated by a pathway from the medial olivocochlear (MOC) system that itself originates in the superior olivary complex (SOC). Spiral ganglion cells making up the auditory-nerve fibers are mainly innervated by connections of the lateral olivocochlear complex (LOC) respectively (Warr and Guinan, 1979; Elgueda and Delano, 2020). Attentional modulations of OAEs can thus be seen as a proxy for subcortical attentional modulations via MOC synapses and have been connected with low-frequency (<10 Hz) oscillatory mechanisms at the cochlear level during alternating selective attention (Dragicevic et al., 2019), with increases in the theta band (∼6 Hz) when attending to upcoming auditory input during an silent cue-target interval (Köhler et al., 2021).

Further evidence for attention modulations of cochlear activity stems from direct recordings in animals with chronically implanted round-window electrodes, showing decreased auditory nerve action potentials to task-irrelevant click sounds during alternate states of visual attention in cats (Oatman, 1971) and during selective attention in chinchillas (Delano et al., 2007). However, whether *human* auditory nerve activity can be directly modulated via selective attention remains unknown.

While direct recordings are normally not feasible in humans, cochlear implants (CI) provide a unique opportunity for recording auditory nerve activity. Besides stimulating nerve fibers inside the cochlea, conventional CI electrodes are used to measure short responses (∼0.2-0.8 ms) to biphasic pulses, so called electrically evoked compound action potentials (ECAPs), to assure auditory nerve and device functioning during and after surgery (Miller et al., 2008; Ramekers et al., 2014). However, the aforementioned effects of selective attention were reflected in slow oscillatory activity <30 Hz that cannot be measured with standard short-latency ECAPs. Assuming as a working hypothesis that selective attention modulates human auditory nerve activity in a similar frequency range, our approach appended short recording windows in a silent cue-target period. This technique allows for discrete sampling of that period within a single trial that can later be processed like standard electroencephalographic (EEG) recordings. Interestingly, CI recipients lack the efferent MOC reflex that leads to cochlear dynamic compression in normal hearing (Wilson et al., 2003; Lopez-Poveda et al., 2016; Marrufo-Pérez et al., 2019). As CI recipients additionally show substantial degeneration of OHCs which are further damaged during surgical insertion of the implant, proposed CI recordings during selective attention should mostly reflect modulations of spiral ganglion cell activity via LOC synaptic connections (Lopez-Poveda, 2018) rather than modulations of scattered OHC populations by residual MOC efferents.

Using an audiovisual crossmodal attention task adapted from Hartmann and Weisz (2019, see **Figure 1A**), we show that ongoing auditory nerve activity in a silent cue-target interval is modulated by focused attention using standard MED-EL CIs as recording devices (see **Figure 1B**). In addition to this average condition-level effect, we show that a classifier is even able to decode attended modality on a single-trial basis, which could have important implications for the use of conventional CIs in a closed-loop system.

**Figure 1.**
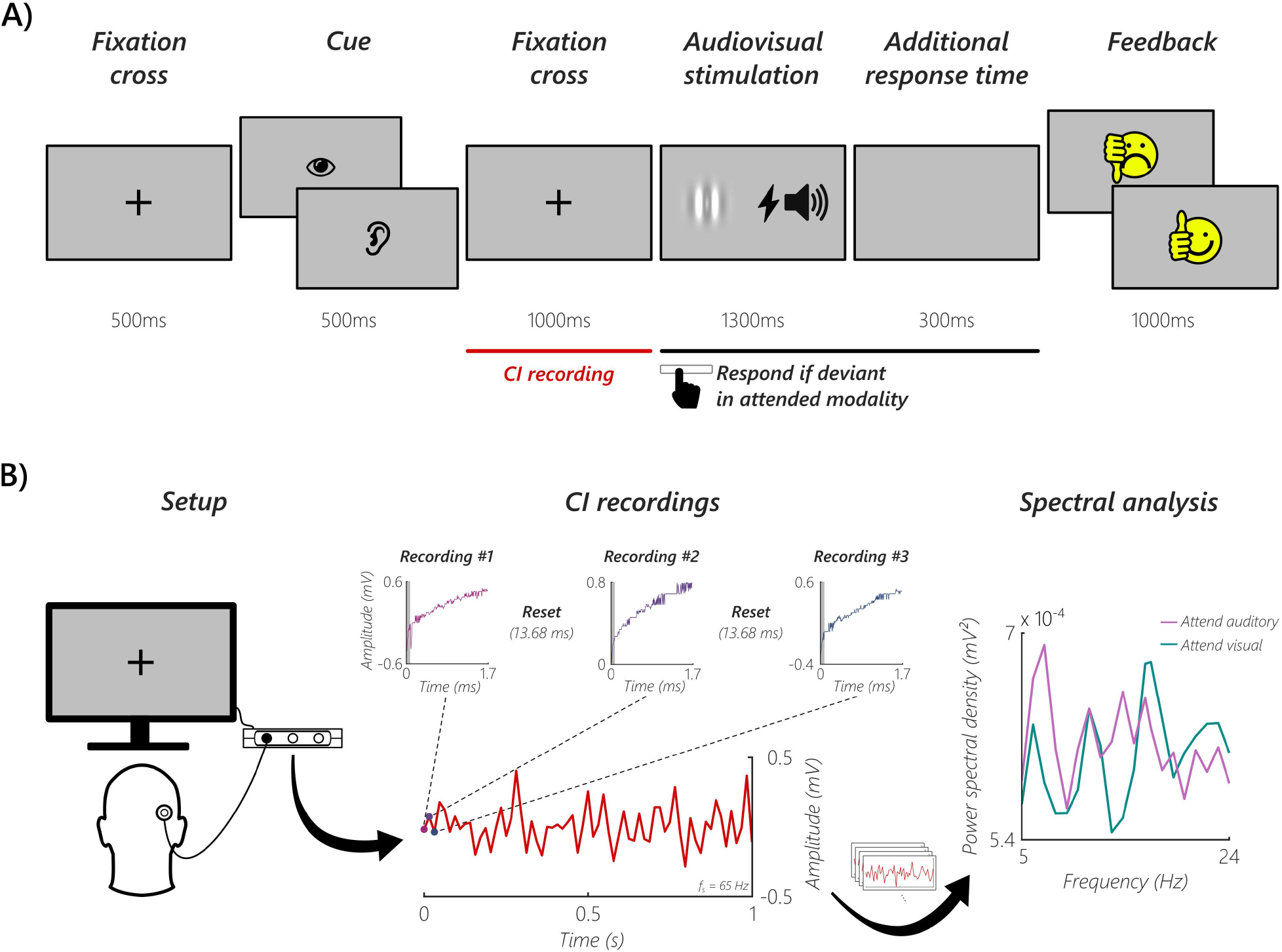
Schematic illustration of the crossmodal attention task and CI recordings. (A) Each trial started with a fixation cross, followed by a cue indicating either to attend the visual or auditory domain. A second fixation cross appeared and an auditory and visual stimulus were presented afterwards. When the stimulus in the attended modality was deviant (visual: gabor patch tilt, auditory: oddball sound), participants had to respond by pressing the spacebar. The additional response time accounted for trials where the gabor patch tilted towards the end of the stimulation. At the end of each trial, feedback was given in the form of a smiley face. The red line denotes the time window where auditory nerve activity was recorded via the CI. (B) Left: Participants were seated in front of a computer screen and were asked to remove their CI processor and coil to replace it with a coil connected to one of the ports of the MAX Programming Interface. Middle: Each recording window was 1.7 ms long, followed by a 13.68 ms reset period (three recordings of an exemplary participant are shown). Due to filter artifacts, the first 100 samples from every recording window were discarded (shaded grey area in the recordings). Each recording was averaged and treated as one sample point. By concatenating these single samples, a recording length of 1 second with a sampling frequency (f_s_) of 65 Hz was reached in every trial. Right: Single trials were averaged and spectral analysis was performed separately for the two conditions.

## Materials and Methods

### Participants

21 right-handed CI users (4 females, *M*_age_ = 57.5, *SD*_age_ = 11.9) participated in the study, all with a minimum CI experience of six months. Participants were recruited via the ear-nose-throat (ENT) departments of the hospitals in Salzburg (n = 10) and Wels-Grieskirchen (n = 11). Three participants were excluded because of a too weak contact between transmitting CI coil and receiver that was required for the study. One participant showed no N1 in recorded ECAPs, which could indicate a measurement problem and was therefore excluded. One participant quit during the session due to concentration problems. This led to a final sample size of 16 participants (3 females, *M*_age_ = 53.8, *SD*_age_ = 12.0). All participants reported no previous neurological or psychiatric disorders, and reported normal or corrected-to-normal vision. All participants signed an informed consent and were reimbursed with 10 Euro per hour. The experimental protocol was approved by the ethics committee of the University of Salzburg and was carried out in accordance with the Declaration of Helsinki.

### Stimuli and experimental design

The experimental procedure was implemented in MATLAB 8.6 (The MathWorks Inc., Natick, Massachusetts, USA) using custom scripts. Presentation of visual stimuli and response collection was achieved with a previous version (th_ptb; https://gitlab.com/thht/th_ptb) of the Objective Psychophysics Toolbox (o_ptb; Hartmann and Weisz, 2020), which adds an additional class-based abstraction layer in addition to the Psychophysics Toolbox (Version 3.0.14; Brainard, 1997; Pelli, 1997; Kleiner et al., 2007). Cochlear stimulation as well as recording was performed via the MAX Programming Interface, a device which is part of the clinical standard setup that enables control of the implant, together with the Research Interface Box 2 Dynamic-link library (RIB2 DLL provided by the University of Innsbruck, Innsbruck, AT; Litovsky et al., 2017). To ensure accurate stimulus presentation and triggering, timings were measured with the Black Box ToolKit v2 (The Black Box ToolKit Ltd., Sheffield, UK).

Participants were seated in front of a computer screen and were asked to remove their CI processor and coil to replace it with a coil connected to the MAX Programming Interface. For bilateral CI users, the side with the better subjective hearing performance and/or longer implantation date was used. Primarily the CI coil model MAX Coil was used, but if the magnet was too weak to ensure a stable connection, the CI coil model MAX Coil S was used. As a first step, the individual electrical hearing threshold was determined with a standard tone with a stimulation frequency of 100 Hz and a duration of 300 ms. To ensure that the auditory stimulation was at a comfortable level during the experiment, the individual maximum loudness was determined, for the standard and an oddball tone respectively. An oddball tone with the maximum possible stimulation frequency of 9990 Hz (based on the used phase duration of 30 µs per phase for sequential biphasic pulses) and a duration of 300 ms was used. The described routines were implemented using custom scripts and the Palamedes Toolbox (Prins and Kingdom, 2018).

Afterwards, as a functionality check of the measurement setup, ECAPs were recorded (Bahmer et al., 2010). ECAPs were biphasic pulses (anodal polarity of the first pulse phase) with a 40 µs phase duration and an 147 µs interpulse interval. In each participant, the first (i.e. most apical) electrode was used for stimulation and the second for recording. Phase amplitudes and amount of ECAPs measured in each participant were defined between the minimum amplitude given by the electrical hearing threshold and the maximum amplitude given by the maximum loudness of the standard tone (phase amplitude: in steps of 9.45 current units (CU); amount: in steps of one).

For the crossmodal attentional task described later, it was necessary that two stimulation frequencies could be distinguished. Because of interindividual differences when hearing with a CI, it was necessary to adjust these stimulation frequencies for every participant. Participants were asked, after hearing a standard and oddball tone, if the first or the second tone had a higher stimulation frequency. The standard tone had a stimulation frequency of 100 Hz and a duration of 300 ms. The initial stimulation frequency of the oddball tone (also with a duration of 300 ms) was determined by the results of the aforementioned maximum loudness procedure. This procedure was carried out using a Bayesian active sampling protocol to estimate the model parameters of the psychometric function (Kontsevich and Tyler, 1999; Sanchez et al., 2016) and was implemented with the VBA Toolbox (Daunizeau et al., 2014). To define the individual oddball stimulation frequency for the subsequent crossmodal attention task, the algorithm searched for the optimal difference in logarithmic steps from 1 to 9890 Hz and this value was subsequently added to the standard stimulation frequency. Six participants heard no clear difference and it was necessary to adjust the oddball stimulation frequency manually, with values between 114 and 600 Hz.

The actual experiment was carried out as a crossmodal attention task (see **Figure 1A**; similar to Hartmann and Weisz, 2019) in six blocks, with 85 trials per block. Each trial started with a 500 ms fixation cross, followed by a cue that indicated either to attend the auditory or the visual modality. Every block had 43 auditory and 42 visual cues. The cue was a picture of an eye or ear, presented for 500 ms. A second fixation cross appeared for 1000 ms and the audiovisual stimulation started afterwards. The auditory stimulation consisted of a 300 ms tone with a stimulation frequency of 100 Hz and was directly presented via the CI coil. The visual stimulation was a vertically oriented gabor patch (spatial frequency: 0.01 cycles/pixel, sigma: 60 pixels, phase: 90°), presented for 1300 ms in the center of the screen. In every block, 8 trials were randomly chosen as visual oddball trials. Independently, another 8 trials were chosen to be auditory oddball trials. Therefore it was possible that a trial was a visual and auditory oddball trial simultaneously. In visual oddball trials, the gabor patch tilted 10° to the left, with a random onset. In auditory oddball trials, a 300 ms tone with the individual oddball stimulation frequency was presented. Participants had to press the spacebar if the current trial had an oddball in the cued domain. To account for trials where the visual oddball onset was towards the end of the stimulation, an additional response time of 300 ms was provided. After each trial, feedback in the form of a smiley face displayed for 1000 ms, indicated if the response was correct or not. To ensure correct understanding of and response during the task, participants completed one block as a practice run before the actual experiment. The total duration of the experiment was about 90 minutes including breaks and preparation.

### Recording of auditory nerve activity

We exploit the ability of CIs to record electrical activity from the cochlea in short time windows, but in contrast to previous approaches (Mc Laughlin et al., 2012; Abbas et al., 2017), in a silent cue-target period. Using a custom developed MATLAB toolbox to abstract MAX Programming Interface commands, we recorded auditory nerve activity via the CI electrode. In every participant, the first (i.e. most apical) electrode was used for the recordings. Each recording window was 1.7 ms long, followed by a 13.68 ms reset period resulting in a sampling frequency of 65 Hz (see **Figure 1B;** 1.7 ms recording + 13.68 ms reset time). The sampling rate of each 1.7 ms recording window was 1.2 Mhz (i.e. 2048 sample points in each recording window). The technical specifics of the measurement system added a random offset to each of the recordings (Gaussian noise, *SD* = 0.4 mV). Because of the USB connection between the computer and the MAX Programming Interface, the start of the first recording window had a jitter of 27 ms, but the system sent a highly precise trigger when it started. Due to technical limitations, it was not possible to record and stimulate simultaneously. We performed recordings in the 1000 ms prestimulus window (see red line in **Figure 1A**).

### Data preprocessing

The raw data was analyzed in MATLAB 9.8 (The MathWorks, Natick, Massachusetts, USA). Due to filter artifacts (using the standard filter from the used RIB2 package), the first 100 samples (= 0.083 ms) from every recording window were discarded. Afterwards, the recording was averaged and treated as one sample point. By repeating these steps for every window and concatenating the single samples, a recording length of 1 second with a sampling frequency of 65 Hz was reached. The data was further preprocessed with the FieldTrip toolbox (revision ea6897bb7; Oostenveld et al., 2010) and a bandpass filter between 4 and 25 Hz was applied (hamming-windowed sinc FIR filter, onepass-zerophase, order: 424, transition width: 0.5 Hz). For one participant, 15 trials had to be rejected because the CI coil fell off during the last trials of one block. Only trials with a correct response were analyzed, which were on average 488 trials (*SD* = 15.8). The number of correct trials was not significantly different between the two conditions (see Behavioral results).

### Frequency analysis

Next, data was demeaned, detrended and power spectral density (PSD) from 4 to 25 Hz was computed on the whole 1000 ms prestimulus window (‘mtmfft’ implementation in FieldTrip with a Hann window) separately for the two conditions. For **Figure 2A**, no bandpass filter was applied, condition-specific power spectra were smoothed (five-point moving average), grand-averaged, and corrected error bars for within-subjects designs were calculated (O’Brien and Cousineau, 2014).

**Figure 2.**
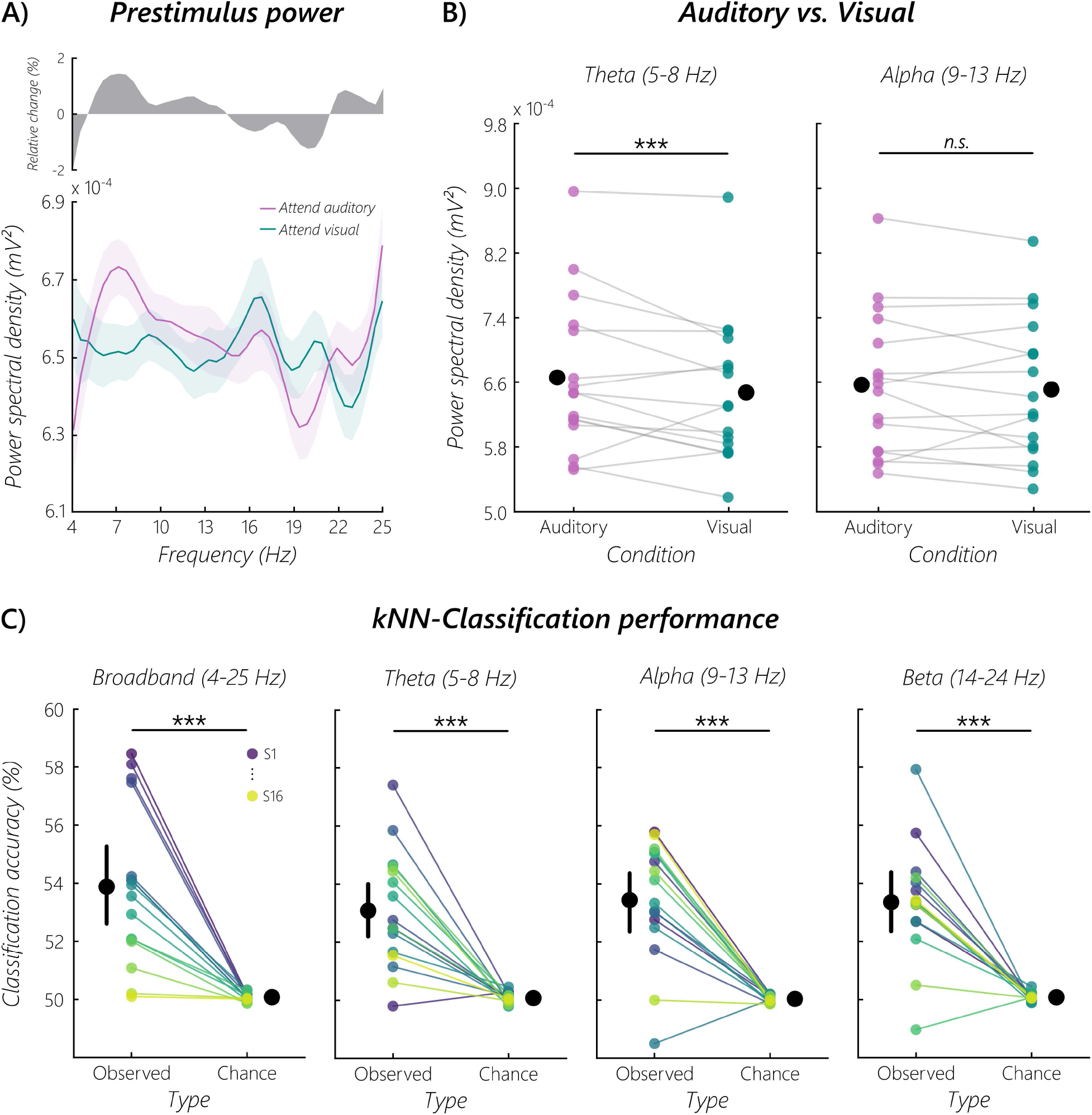
Prestimulus power modulations and decoding of selective attention. (A) Grand average prestimulus power spectra from 4-25 Hz when attending the auditory or visual domain. The top panel indicates the relative change between the auditory and visual domain. The shaded areas in the bottom panel represent the standard error of the mean for within-subjects designs (O’Brien and Cousineau, 2014). (B) Average prestimulus power in the theta and alpha band, separated by the two conditions. Black dots indicate the group mean for the respective condition. A cluster-based permutation test in the averaged theta FOI resulted in a statistically significant difference when testing the hypothesis that performance is higher when attending the auditory domain (*p* = 10.00e^-05^, *d* = 0.49). No cluster was found in the alpha band (*p* = 1.00, *d* = 0.20). The asterisks indicate a statistically significant difference (n.s. = not significant). (C) A kNN-Classifier was used to decode attended modality from single-trial prestimulus power spectra. Resulting *Observed* accuracies were contrasted with respective *Chance* levels of a random permutation test for all FOI. Contrasts revealed significant (*p* < 0.001) decoding performance throughout spectra with fairly similar effects (broadband: *t*(15) = 5.60, *p* = 2.60e^-05^, *d* = 1.96; theta: *t*(15) = 5.83, *p* = 1.70e^-05^, *d* = 2.11; alpha: *t*(15) = 6.78, *p* = 3.00e^-06^, *d* = 2.34; beta: *t*(15) = 6.40, *p* = 6.00e^-06^, *d* = 2.33) on a group level (represented by black dots; error bars = 95% CI). However, on a single-subject level the attention effect was most pronounced for individually specific FOI, resulting in significant above chance decoding for 12 out of 16 subjects. As indicated in the top right of the broadband column (S1: purple, S16: yellow), each subject is represented by the same color in all four FOI columns.

We defined two frequency bands of interest (FOI), theta (5-8 Hz) and alpha (9-13 Hz). Theta was selected because of previous work on OAEs that showed attentional modulations in this FOI (Dragicevic et al., 2019; Köhler et al., in press). On a cortical level, previous work showed that auditory alpha activity reflects attentional processes (Weisz et al., 2011, 2014; Müller and Weisz, 2012; Frey et al., 2014; Mazaheri et al., 2014; Weise et al., 2016). Therefore, we decided to analyze this FOI at the cochlear level.

### Decoding analysis

For decoding of attended modality on a single-trial basis, we performed k-nearest neighbors (kNN) classification of single-trial power spectra using scikit-learn (Version 0.23.1 running on Python 3.7.7; Pedregosa et al., 2011) separately for a broadband signal (4-25 Hz) followed by standard frequency bands associated with selective attention (theta: 5-8 Hz, alpha: 9-13 Hz, beta: 14-24 Hz). We decided to use the kNN classification approach as data was recorded from a single CI channel over a one second period resulting in low numbers of features (i.e. frequency points per band), a classification problem usually solved better by a kNN approach (Eisa et al., 2018).

At first, a subject’s data was standardized to unit variance and zero mean. For the classification process of each subject, the best number of neighbors was determined by searching the hyper-parameter space for the best cross-validation (CV) score of a kNN model using the implemented *GridSearchCV* function with a 2-fold CV on shuffled class samples (*StratifiedKFold(shuffle=True)*) that was fit to the data for every FOI. Our decision for a 2-fold CV was based on recommendations in case of low sample / effect size data (Jamalabadi et al., 2016). The numbers of neighbors to use during the gridsearch were defined as ranging from one to 10% of trials in the dataset in odd numbers (1, trials/10, stepsize=2) to avoid the conflict of even neighbors in a two-class problem (attend auditory vs. visual).

### Statistical analysis

#### Cluster-based permutation statistics of the power spectra

To test the hypothesis that power was higher when the auditory domain was attended, statistical testing of PSD was performed with a cluster-based permutation test (dependent samples t-test, 10000 randomizations, one-tailed; Maris and Oostenveld, 2007). We averaged the theta and alpha FOI and tested them separately.

#### Decoding analysis statistics

Given the novel approach, we could not exclude that the classifier would pick up on a few outlying data points. In order to address this issue explicitly, the classifier was tested on the same noisy data, albeit with randomly shuffled condition labels. Samples were thus classified and tested for significance with the best scoring number of neighbors in a 1000 random permutation test and the aforementioned 2-fold CV procedure. The resulting Observed and Chance accuracy values (where chance level was calculated as the mean accuracy of the 1000 random permutation scores) for every FOI were then statistically tested using pingouin (Version 0.3.8 running on Python 3.7.7; Vallat, 2018). In a first step, to test whether auditory nerve modulation was generally reflected within classification results, broadband values (Observed vs. Chance) were compared using a one-sided t-test. Then, classification results of all four FOI were compared in a two-factor repeated measures ANOVA with the factors FOI (broadband, theta, alpha, beta) and Type (Observed vs. Chance) to check whether the attention effect was driven by one of the predefined FOI. Finally, theta, alpha, and beta bands were also tested for significant differences during focused attention computing three one-sided t-tests with respective values (Observed vs. Chance).

### Code Accessibility

The data and code necessary for statistical analysis and generating the figures are available at the corresponding author’s GitLab repository (https://gitlab.com/qubitron).

## Results

Sixteen CI users performed a crossmodal attention task (similar to Hartmann and Weisz, 2019) where attention had to be focused on an upcoming auditory or visual stimulus (see **Figure 1A**). Auditory nerve activity was recorded directly via their first (i.e. most apical) CI electrode in the silent cue-target interval. We calculated the power spectral density of the signal and compared the two conditions (attend auditory vs. visual) in the theta and alpha band. Afterwards, a classifier was utilized to decode the attended modality on a single-trial basis using the broadband signal and frequency bands typically associated with selective attention (theta, alpha, beta).

### Behavioral results

Participants gave a correct response in 96% (*SD* = 2.7%) of all trials. The number of correct trials did not differ significantly between the two conditions, according to a dependent sample t-test (auditory: *M* = 245 (*SD* = 9.8); visual: *M* = 242 (*SD* = 8.8); *t*(15) = 1.32, *p* = 0.21, *d* = 0.33). When there was an oddball in the cued domain, a correct response was given in 75% (*SD* = 19.0%) of the trials. In the auditory condition, the percentage of correct oddball trials was 72% (*SD* = 30.8%) and in the visual condition 78% (*SD* = 14.2%). Overall, the behavioral findings suggested that participants performed the task in a compliant manner.

### Human auditory nerve activity is modulated by selective attention

In a first analysis step, we calculated the broadband PSD from 4-25 Hz, separately for each condition (attend auditory/attend visual). The resulting power spectra (see **Figure 2A**) by themselves showed no clear peaks, however the grand average condition contrast spectrum indicates differences that are mainly centered in distinct frequency ranges. Based on previous OAE and M/EEG work (Mazaheri et al., 2014; Köhler et al., 2021), we statistically compared the two conditions in the theta (averaged between 5-8 Hz) and alpha frequency band (averaged between 9-13 Hz; **Figure 2B**). A cluster-based permutation test in the theta frequency band showed that prestimulus power is higher when attending the auditory domain (*p* = 10.00e^-05^, *d* = 0.49). No cluster was found in the alpha frequency band (*p* = 1.00, *d* = 0.20). Given the distribution of the individual average power in both FOI, a rather high interindividual variability can be seen. In addition, we also tested the beta frequency band (averaged between 14-24 Hz), no cluster was found (*p* = 1.00, *d* = -0.16).

We further wanted to investigate whether the effect in the theta band was related to behavioral performance. Correlating individual average theta power in each condition with behavioral performance showed a very small and non-significant relationship (auditory: *r* = 0.019, *p* = 0.944; visual: *r* = 0.096, *p* = 0.723). A potential explanation could be the rare occurrence of oddball trials (see Materials and Methods), resulting in a very high overall behavioral performance (auditory: *M* = 96%, *SD* = 3.1%; visual: *M* = 95%, *SD* = 3.6%).

The results so far showed that selective attention modulates directly recorded cochlear activity, with the effect being in particular pronounced in the theta frequency range: attending to an upcoming auditory stimulus resulted in higher power recorded from the CI electrode.

### Auditory nerve as origin of the signal

Our results so far suggest a theta rhythmic modulation of the human auditory nerve by selective attention, with increased theta activity when attending an upcoming auditory stimulus. However, one concern of the recording approach was the origin of the recorded signal being actually any cortical source instead of auditory nerve activity. Even though electrode configurations with close (extracochlear) reference on the implant housing should already pick up quite local sources, an empirical approach was needed to exclude possible effects of volume conduction. To show that our effects truly resemble auditory nerve modulations, we exploited an additional EEG dataset that was simultaneously recorded to the CI signal for one of the participants. Addressing this issue, we cross-correlated the CI signal with the EEG signals of all 64 electrodes. If signals at the CI were due to volume conduction, this should go along with strong cross-correlations at zero-lag. To test the absence of this instantaneous volume conduction effect, we calculated the Bayesian Pearson correlation between CI and EEG signals. We report ln-transformed Bayes factors, ln(BF_10_), where values > 1.1 would mean substantial evidence for volume conduction and values > 2.3 would mean strong evidence for volume conduction (Jeffreys, 1998). Results of this analysis show no evidence for volume conduction according to ln(BF_10_): *M* = -0.98 (*SD* = 0.40), with values ranging from -1.60 to 0.20 between all electrodes. The analysis substantially strengthens our argument that the attentional effects are genuinely recorded from the auditory nerve.

### Attended modality can be decoded from single-trial CI recordings

We used prestimulus power spectra for a kNN-Classifier to show that attention modulation of ongoing auditory nerve activity in humans is even reflected in single-trial CI recordings. To ensure that the classifier was able to differentiate auditory nerve activity when attending the auditory compared to the visual domain in general, we calculated a t-test between *Observed* classification accuracies and respective *Chance* levels of broadband power spectra, showing that this attention effect was decodable significantly above chance (*t*(15) = 5.60, *p* = 2.60e^-05^, *d* = 1.96; **Figure 2C**). Given the significant difference over a broad frequency range, we were further interested in whether this attention effect was driven by one of the FOI usually connected with selective attention in OAE and M/EEG studies. We therefore calculated a two-factor repeated measures ANOVA to compare the effect of selective attention on kNN-Classification accuracy for different FOI (broadband, theta, alpha, beta) and Types (*Observed, Chance*). Results show no significant effect of FOI (*F*(3, 45) = 0.37, *p* = 0.78, η_p_^2^ = 0.02), yet show a significant effect for *Observed* vs. *Chance* accuracies (*F*(1, 15) = 136.55, *p* = 6.21e^-09^, η_p_^2^ = 0.90), with higher accuracies for *Observed* (*M* = 0.53) than *Chance* (*M* = 0.50) levels. No FOI x Type interaction on decoding results was found (*F*(3, 45) = 0.36, *p* = 0.78, η_p_^2^ = 0.02). As a main effect of Type and no interaction of FOI x Type indicated that selective attention can be decoded from all FOI separately, in addition we computed three t-tests for theta, alpha, and beta bands contrasting respective *Observed* and *Chance* levels. For all three FOI a significant difference was found for selective attention decoding (theta: *t*(15) = 5.83, *p* = 1.70e^-05^, *d* = 2.11; alpha: *t*(15) = 6.78, *p* = 3.00e^-06^, *d* = 2.34; beta: *t*(15) = 6.40, *p* = 6.00e^-06^, *d* = 2.33; **Figure 2C**). Additionally, random permutation tests of kNN classification within all four different FOI gave insights into single-subject decoding performance across the different frequency spectra. Independent of FOI, an overall number of 12 subjects (i.e. 75% of the sample) showed significant (*p* < 0.05) above chance decoding of focused attention during the silent cue-target interval.

## Discussion

The efferent auditory system comprises a complex arrangement of subcortical pathways, which can alter cochlear activity by top-down signals (Terreros and Delano, 2015; Elgueda and Delano, 2020). Profound evidence supports the notion of altered oscillatory neural activity by selective attention on a cortical level within the alpha (Frey et al., 2014; Mazaheri et al., 2014; Weise et al., 2016) and beta band (Buschman and Miller, 2007; Iversen et al., 2009; Lee et al., 2013). Much less is known for subcortical structures along the efferent pathway, especially when it comes to the *human* cochlea as special recording and analysis techniques are required (Elgueda and Delano, 2020). So far, investigating attentional modulation of cochlear activity in humans had to rely on indirect recordings of OAEs, a noninvasive approach for measuring OHC activity. Recent evidence suggests slow modulations (<10 Hz) of cochlear activity (Dragicevic et al., 2019) that is even enhanced during a silent cue-target period when attending the auditory modality (Köhler et al., 2021). However, studying OAEs cannot address direct modulation of auditory nerve activity since spiral ganglion cells are efferently innervated by a separate, lateral pathway. In contrast to the MOC, most LOC fibers project to the ipsilateral cochlea (Robertson, 1985) and are evenly distributed from apical to basal end (Guinan, 1996). In humans, little is known about their function in sound processing since its unmyelinated axons are difficult to electrically stimulate and record from (Guinan, 2018), while measured responses are inconclusive about MOC or LOC origin. To our knowledge, attentional modulation via the LOC remains completely unknown as direct recordings of auditory nerve activity are normally not feasible in humans. Given the absence of the efferent MOC reflex in CI recipients (Wilson et al., 2003; Lopez-Poveda et al., 2016; Marrufo-Pérez et al., 2019) in addition to substantial OHC degeneration, potential alterations of respective auditory nerve activity in a selective attention paradigm should largely reflect top-down signals from the LOC (Lopez-Poveda, 2018). Our results show that ongoing auditory nerve activity is top-down modulated, putatively suggesting a role of the LOC pathway in selective attention. Future applications that are able to simultaneously record from multiple CI electrodes could exploit the different electrode positions along the cochlea and draw conclusions about frequency specific terminal distributions of LOC efferents. Setups with bilateral CI recordings could even address the LOC’s role in analysis of interaural differences in frequency and intensity, as assumed by Ciuman (2010).

Importantly, we ruled out any cortical source to drive the demonstrated effects as Somers et al. (2021) recently showed that tailored CIs can be used to intentionally obtain evoked potentials from the auditory cortex with a recording setup that is fairly comparable with only small differences to the one used in this study. Whilst Somers et al. used reference electrodes located on the temporal muscle, MED-EL implants have reference and ground electrodes in the actual implant housing. It was therefore important to rule out any cortical origin with additional analysis. Cross-correlation of simultaneously recorded CI and EEG electrodes for one participant clearly showed the absence of any instantaneous correlations which would be caused by volume conduction.

While cortical and OAE-based measures suggest attention-related effects in distinct frequency bands (Mazaheri et al., 2014; Köhler et al., 2021), our results are mixed in this respect. The broadband frequency analysis of the prestimulus interval showed no clear peaks (**Figure 2A**). This, however, may also be the result of low signal-to-noise ratios (SNRs), as commercial CIs are so far not optimized to do these kinds of continuous electrophysiological recordings. Indeed a grand average of the condition differences points to maximal effects in a frequency range overlapping with the one reported by Köhler et al. (2021), resulting in enhanced theta power while attending to the auditory modality (**Figure 2B**). This result corroborates our previous finding using a similar paradigm, where ongoing OAEs in the theta band (∼6 Hz) were enhanced while attending an upcoming auditory stimulus (Köhler et al., 2021). We found no selective attention effect in the alpha nor in the beta band in concordance with aforementioned studies of otoacoustic activity. It is therefore possible that these frequency bands do not play a central role in selective attention at the peripheral level. Further studies with an optimized recording setup will be necessary to address this issue. However, the dominance of theta rhythmic modulations in the auditory periphery across the reported studies are striking. Arguably, feature selection on a cochlear level shares top-down mechanisms with working memory (WM) prioritization processes reflected within the theta band (Riddle et al., 2020). In this context, selective attention could support WM processes in its ability to exclude irrelevant information, i.e. WM filtering efficiency (Arnell and Stubitz, 2010) or expectation-driven WM benefits (Bollinger et al., 2010). This could explain our results with regards to predominance of theta oscillations. However, we think that condition related increases in the theta band in the current study mainly reflect processes of selective attention rather than visual WM since the cues were fairly similar and the task not demanding in terms of WM capacity. This idea should be further explored in future studies that additionally manipulate memory load and assess WM performance outcomes and how this is affected by individual theta in-/decreases already at a cochlear level.

Building upon conventional analyses of condition-level fast Fourier transform (FFT) averages, we decided to use single-trial frequency spectra to classify anticipatory attentional focus during the silent cue-target period. With this approach we aimed to get more detailed insight into fine-grained differences between attentional states coded within modulations of direct cochlear recordings that could be missed by condition-level averaging approaches and indirect OAE measurements. Strikingly, classification of the broadband signal (4-25 Hz) revealed significantly improved differentiation of attended modality compared to the average condition-level effect of the FFT results (**Figure 2C**). Follow-up analysis showed that the performance was not driven by one of three FOI (theta, alpha, beta) usually associated with selective attention, but instead it revealed that the contribution of each of these frequency bands to broadband classification was fairly similar. However, the decoding approach allowed for additional insight into single-subject classification performance and showed high interindividual variability in terms of an optimal spectral frequency band. It remains to be determined whether this effect is driven by local idiosyncrasies at the peripheral level (e.g. synaptic connections between LOC and spiral ganglion cells) or even involves particular activity patterns at higher hierarchical levels. Independent of the precise origins of our effects observed at the auditory nerve, the decoding results open up avenues to future developments towards closed-loop CIs that incorporate mental states of the recipient into adaptive stimulation in real time. As we show, a classifier could use the frequency information of this signal to anticipate the attentional state of the recipient. Future research will need to address which cognitive states can be decoded directly at the auditory nerve and how this information could be exploited in a closed-loop CI setup. In this regard, overall classification results might seem rather small with accuracies ranging between 50% - 60%, yet they are comparable to many others in the field of cognitive neuroscience. Considering that the results are based on simplistic decoding of noisy single-channel data, the reported accuracies are highly valuable and most importantly interpretable. More complex algorithms (e.g. deep neural networks) would most probably lead to much higher decoding accuracies, however they would not add significantly to increase interpretability. Future approaches with improved recording setups would most probably allow for higher decoding accuracies as they were necessary for functional closed-loop systems. Additionally, albeit substantial interindividual variability the classifier yielded accuracies at a quite consistent above chance level. It remains an open question to what extent interindividual differences are functionally relevant. In the current study, we did not observe any relation to behavioral performance, however this may be due to the simplistic task leading to a ceiling effect obscuring associations.

In summary, this study shows that individuals with a CI form a model population to deepen our understanding of how cognition can lever the efferent auditory system to modulate auditory input at the earliest stages of processing. To the best of our knowledge our study is the first to investigate attentional effects on activity recorded directly from the auditory nerve in humans. We confirm and extend previous indirect measurements, suggesting attentional modulations in the theta frequency range. Importantly, we also show that selective attention can be decoded above chance at a single-trial and even individual level. Previous reports on attentional modulations of cochlear activity relied on OAEs, which are driven by the MOC pathway. Our results strongly suggest that the LOC pathway can also be exploited in a top-down fashion to affect spiral ganglion cells directly.

## Acknowledgments

Q.G. and P.R. are supported by the Austrian Research Promotion Agency (FFG; BRIDGE 1 project “SmartCIs”; 871232) and the Austrian Science Fund (FWF; Doctoral College “Imaging the Mind”; W 1233-B). The authors would like to thank Agnes Koller and Lisa Niederwanger for their help in recruiting participants.

